# *Pseudomonas aeruginosa* detachment from surfaces via a self-made small molecule

**DOI:** 10.1101/2020.07.14.203174

**Authors:** Robert J. Scheffler, Yuki Sugimoto, Benjamin P. Bratton, Courtney K. Ellison, Matthias D. Koch, Mohamed S. Donia, Zemer Gitai

## Abstract

*Pseudomonas aeruginosa* is a significant threat in both healthcare and industrial biofouling. Surface attachment of *P. aeruginosa* is particularly problematic as surface association induces virulence and biofilm formation, which hamper later antibiotic treatments. Previous efforts have searched for biofilm dispersal agents, but there are no known factors that specifically disperse surface-attached *P. aeruginosa*. In this study we develop a quantitative surface-dispersal assay and use it to show that *P. aeruginosa* itself produces factors that can stimulate its dispersal. Through bioactivity-guided fractionation, Mass Spectrometry, and Nuclear Magnetic Resonance, we elucidated the structure of one such factor, 2-methyl-4-hydroxyquinoline (MHQ). MHQ is an alkyl-quinolone with a previously unknown activity and is synthesized by the PqsABC enzymes. Pure MHQ is sufficient to disperse *P. aeruginosa*, but the dispersal activity of natural *P. aeruginosa* conditioned media requires additional factors. Whereas other alkyl quinolones have been shown to act as antibiotics or membrane depolarizers, MHQ lacks these activities and known antibiotics do not induce dispersal. In contrast, we show that MHQ inhibits the activity of Type IV Pili (TFP) and that TFP targeting can explain its dispersal activity. Our work thus identifies surface dispersal as a new activity of *P. aeruginosa-*produced small molecules, characterizes MHQ as a promising dispersal agent, and establishes TFP inhibition as a viable mechanism for *P. aeruginosa* dispersal.

**Significance Statement:** We discovered that the clinically relevant human bacterial pathogen *P. aeruginosa*, typically associated with surface-based infections, is dispersed by a small molecule that the bacteria themselves produce. We elucidate the chemical structure of this molecule and find that mechanistically it functions to inhibit the activity of the *P. aeruginosa* extra cellular surface motility appendage, the type IV pilus.

## Introduction

Hospital-acquired infections (HAIs) and the pathogens that cause them are a growing concern due to the rise in antibiotic resistance, which makes treating these infections increasingly difficult (1). Multi-drug-resistant pathogens are particularly problematic in health care settings as hospital patients are often immunocompromised and contaminated surfaces promote bacterial transfer to new patients. The list of contaminated surfaces ranges from medical implants to neckties worn by doctors.

One of the major causes of a wide variety of HAI’s is the bacterium *Pseudomonas aeruginosa* (1, 2). *P. aeruginosa* inhabits a wide range of environments and can infect a surprising array of hosts due to its large number of virulence factors(3, 4). Recently, our lab and others demonstrated that surface attachment strongly induces *P. aeruginosa* virulence (5–7). To initiate surface-induced virulence, *P. aeruginosa* senses surfaces through type IV pili (TFP), which are extracellular polymers that can be actively extended and retracted. Thus, disrupting surface attachment or TFP activity could be powerful yet largely untapped methods to combat *P. aeruginosa* pathogenesis, by reducing its propensity to disseminate via surfaces and by reducing the induction of its virulence mechanisms.

We sought to identify new ways of disrupting surface attachment through chemical means such as small molecules. *P*. *aeruginosa* produces many excreted metabolites with a wide array of functions (8). These secreted factors include virulence factors such as pyocyanin and rhamnolipids, as well as factors that regulate specific aspects of the *P. aeruginosa* life cycle like the biofilm dispersal cue, *cis*-2-decenoic acid (9). A number of groups have screened for compounds that disrupt biofilms (9–12), but significantly less work has been done on inhibitors of surface attachment itself.

Here we show that cell-free supernatant from cultures of *P. aeruginosa* PA14 cause rapid dispersal of cells from the surface. Using a bioactivity-guided chemical fractionation and purification approach, we identify 2-methyl-4-hydroxyquinoline (MHQ) as a small molecule factor made by *P. aeruginosa* that induces its own dispersal. MHQ has not been previously characterized with respect to its function or synthesis. We show that MHQ is synthesized by enzymes in the *Pseudomonas* quinolone signaling (PQS) pathway. Furthermore, we find that MHQ inhibits TFP pilus activity, potentially explaining how MHQ causes dispersal of *P. aeruginosa* from the surface. Our findings thus characterize a previously understudied secreted metabolite with the potential to prevent *P. aeruginosa* infections by limiting surface attachment.

## Results

### P. aeruginosa *conditioned media removes* P. aeruginosa *from a surface*

Since *P. aeruginosa* is known to produce a staggering array of secreted secondary metabolites with a variety of biological functions, we investigated whether it might produce a compound that would cause cells to detach from a surface. To test a variety of conditions in a high throughput manner we designed an assay we term DISPEL, for <u>d</u>ispersal of <u>i</u>nitially <u>s</u>urface-attached <u>p</u>athogens via <u>e</u>xtract <u>l</u>avage. In brief, *P. aeruginosa* cells were first allowed to attach to the surface of a 96 well plate, the unattached cells were removed by washing, the attached cells were treated with molecules or supernatants of interest, the cells were then washed to remove dispersed cells, and the plate was imaged to determine the number of surface-attached cells remaining (Fig. 1A-B). Figure 1C shows a quantification of the number of mid-log recipient cells that remained attached after treatment with phosphate buffered saline (PBS), Luria–Bertani medium (LB), or cell free supernatant from an overnight culture.

**Figure 1.**
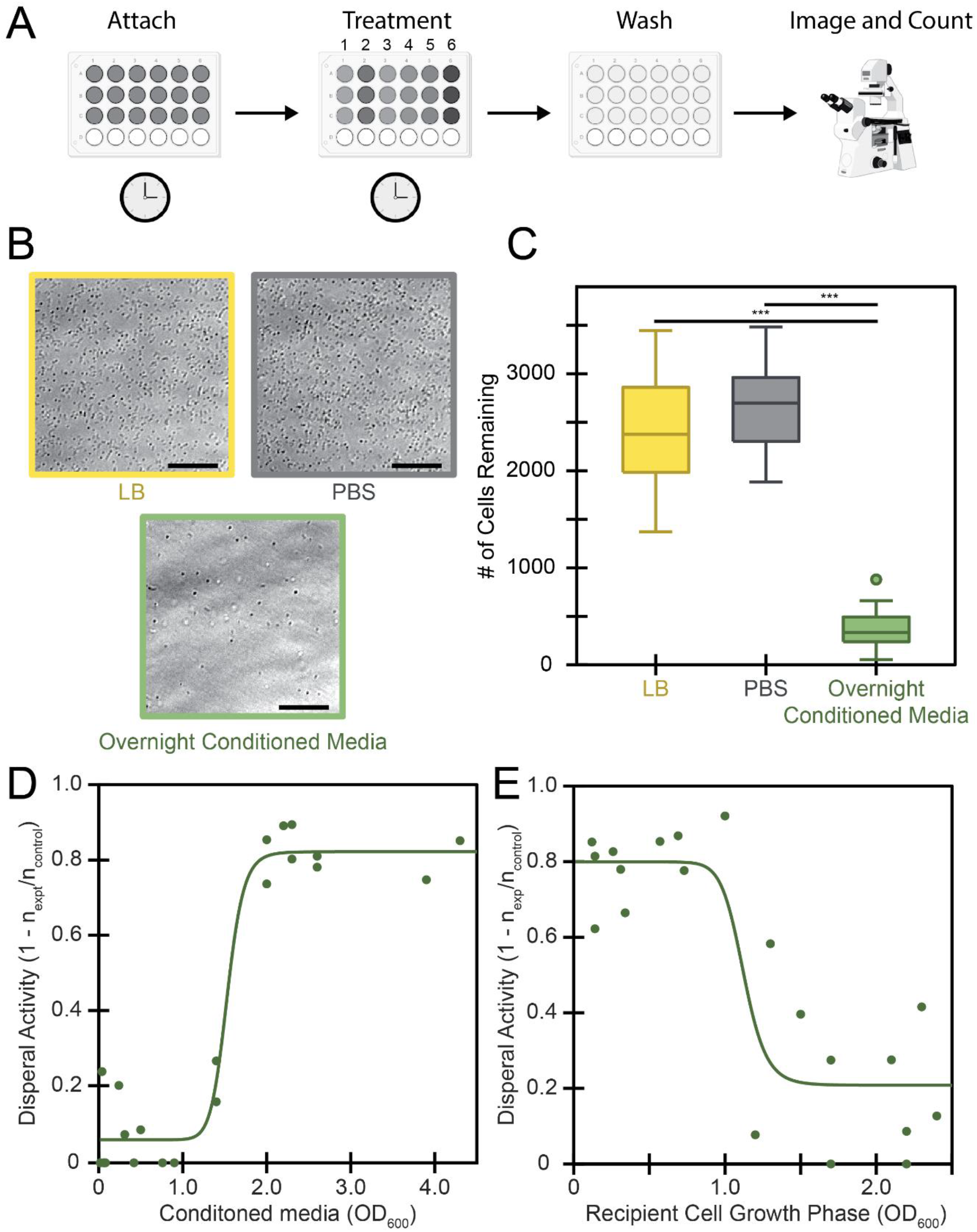
Conditioned media from *P. aeruginosa* disperses *P. aeruginosa* cells. (A) Schematic of the DISPEL assay. <u>D</u>ispersal of <u>i</u>nitially <u>s</u>urface-attached <u>p</u>athogens via <u>e</u>xtract <u>l</u>avage (DISPEL) involves the following: attaching cells for 10 minutes; removing planktonic culture; treating attached cells for 10 minutes; gently washing to remove treatment and detached cells; imaging and counting remaining attached single cells. (B-C) Control treatments of LB or PBS are compared to experimental treatments such as cell free media from an overnight culture. (B) Example phase contrast micrographs (scale bar 40 μm). (C) Quantification of micrographs depicted in box and whisker plots from 24 biological replicates. *** p-value < 0.001 from Student’s t-test. (D-E) Dispersal activity is bounded between 0 (no cells removed) and 1 (all cells removed) and is calculated as 1 – (cells still attached post experimental treatment / cells still attached post control treatment). Fits are based on modified Hill equation (Equation S2). (D) Dispersal activity of cell free conditioned media from varied OD_600_ cultures on mid-log (OD_600_ 0.6-0.8) *P. aeruginosa* cells. (E) Dispersal activity of cell free overnight conditioned media on *P. aeruginosa* cells from varied OD_600_ cultures. Before attachment cultures were diluted or concentrated to standardize the seeding density.

To determine when *P. aeruginosa* produces a dispersal signal, we used the DISPEL assay to test the dispersal activity of cells from different growth phases. We therefore tested cell free supernatant from *P. aeruginosa* cultures grown to different densities for their ability to disperse mid-log *P. aeruginosa* (OD = 0.6-0.8) (Fig. 1D). We defined dispersal activity as the fraction of cells removed by the treatment (1 − n_(cells remaining after treatment)_ / n_(cells remaining after control)_). As controls, cells treated with either LB growth medium or PBS remained attached to the surface in large numbers with little to no dispersal activity relative to one another (Fig. 1B-C). In contrast, cells treated with cell free overnight conditioned media and with supernatants from cultures of OD_600_ greater than 1.5 were largely removed from the surface (Fig. 1D). Conditioned media from cultures at OD_600_ less than 1.5 showed little to no activity (Fig. 1D).

Because dispersal activity of supernatant is dependent on the culture growth phase, we also sought to determine whether dispersal response of the attached cells is growth phase specific. To test this, we grew *P. aeruginosa* over a range of cell densities, normalized their cell numbers, and allowed those cells to attach to the surface. We then treated each sample with conditioned media from late stationary growth phase cultures of *P. aeruginosa* and quantified the extent of dispersal. Recipient cells that were below an OD_600_=1.0 showed high dispersal whereas cells above OD_600_=1.0 showed low dispersal (Fig. 1E). These data show that as *P. aeruginosa* cultures become denser, their supernatants increase in dispersal activity, but the cells themselves become less responsive to the dispersal cue, suggesting a mechanism for growth-phase specific dispersal and attachment. To further study this phenomenon, we thus focused on overnight-grown signalers and mid-log recipients.

### Bioactivity-guided fractionation identifies MHQ as a *P. aeruginosa* dispersal agent

To characterize the chemical nature of the dispersal activity in *P. aeruginosa* supernatants we first used a variety of perturbations. These results suggested that the activity is mediated by a small organic molecule as the activity was protease and nuclease insensitive, heat stable, and < 5 kDa in size. Consequently, we generated 50L of PA14 cell-free supernatant from overnight cultures and isolated the small molecules using Diaion HP-20 resin followed by ethyl acetate extraction. This crude extract was fractionated on an open Mega Bond Elut-C18 column. Two of these fractions retained significant dispersal activity, confirming the initial indications that the activity is mediated by a small molecule (Fig. 2A).

**Figure 2.**
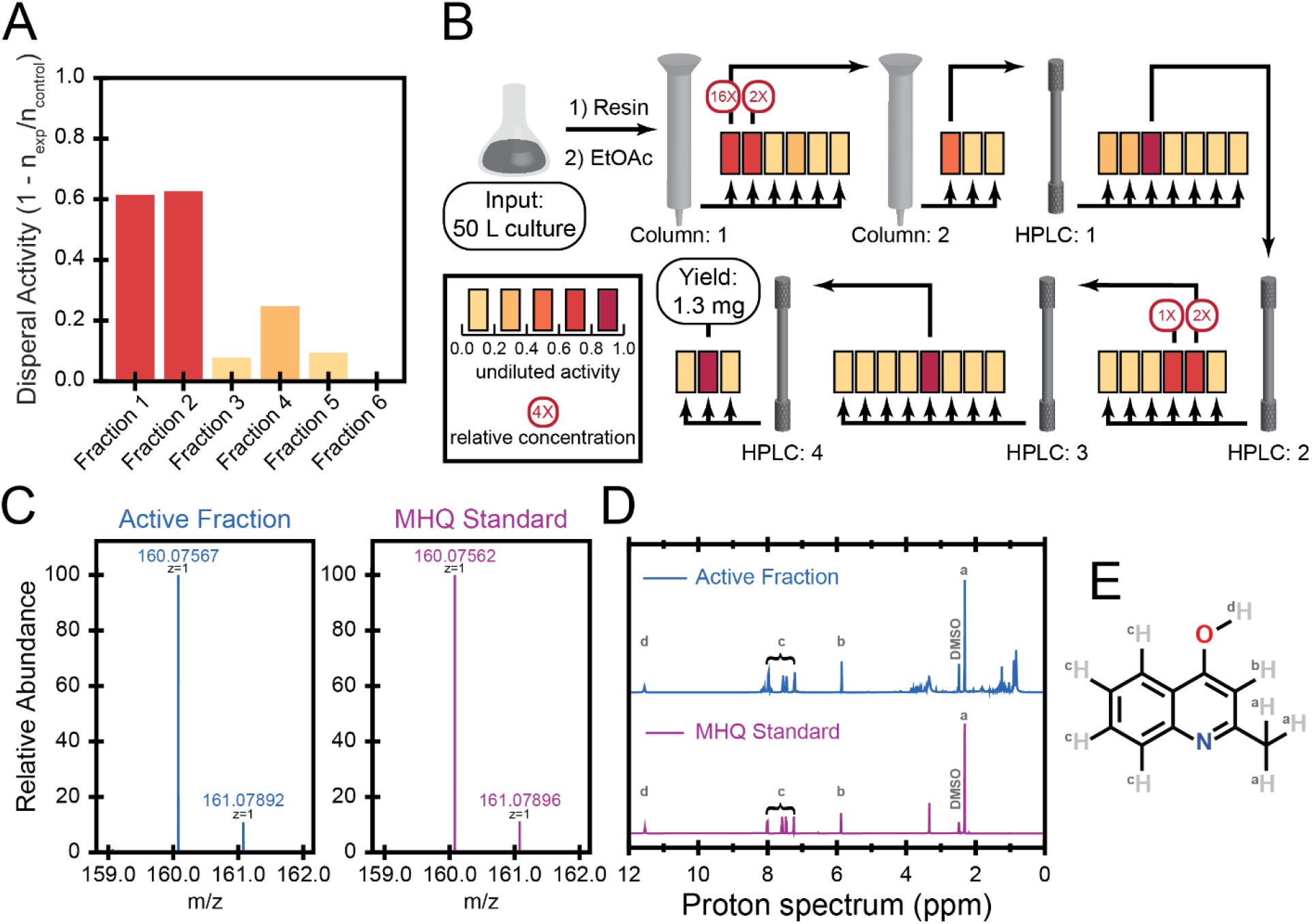
Bioactivity-guided fractionation identified MHQ as a *P. aeruginosa* dispersal agent. (A) Fractionation of crude extract from 50L of overnight conditioned media of *P. aeruginosa* revealed that the first two fractions contained dispersal activity against mid-log (OD_600_ 0.6-0.8) *P. aeruginosa* cells. (B) We utilized our DISPEL assay to perform bioactivity-guided fractionation. Multiple rounds of fractionation continued to retain activity until the sample was pure enough for further chemical analysis. (C) High-resolution mass spectrum of the purified MHQ in the final active fraction (**VI**). (D) ^1^H-NMR spectrum of purified MHQ in the final active fraction **VI** in comparison to that of an authentic standard of the same molecule obtained commercially. See also Fig S2-4 and Table S3 for additional NMR spectra and peak assignments. (E) MHQ is an alkyl-quinolone with a single methyl as the tail group.

To determine which fraction had a higher concentration of active molecule we assayed serial dilutions of the active fractions and determined that the first fraction (**I**) had ~8-fold higher concentration of active molecule. We thus sub-fractionated **I** using a second open Mega Bond Elut-C18 column and identified a single sub-fraction (**II**) that retained dispersal activity. **II** was further sub-fractionated using four subsequent rounds of High Performance Liquid Chromatography (HPLC) C-18 column fractionation (we designated the sub-fraction with the highest activity in each round with roman numerals **III-VI**). Activity of the extract was assessed after each round of fractionation to ensure activity was not lost throughout the process (Fig. 2B).

The final round of fractionation yielded 1.3 mg of total material (**VI**) that was active in the dispersal assay. HPLC coupled with Mass Spectrometry (HPLC-MS) and HPLC high resolution-MS (HPLC-HRMS) analyses of this final sub-fraction indicated that it was dominated by a single molecular species with a mass-to-charge ratio (m/z) of 160.076 [M-H]^+^ (Fig. 2C), which corresponds to a molecule with a predicted mass of 159.068. From this mass we determined the most likely chemical formula of the dominant species as C_10_H_9_NO. To determine its structure, we performed Nuclear Magnetic Resonance (NMR) analysis on **VI**, which revealed the dominant molecule’s structure to be 2-methyl-4-hydroxyquinoline (MHQ) (Fig. 2D-E, Fig. S1-3, Table S3). Finally, we obtained a commercial standard of MHQ (Sigma-Aldrich) and confirmed that it matched the active fraction exactly with respect to its retention time, MS, and NMR, thereby validating the structural elucidation (Fig. 2C-D, Fig. S1-4, Table S3). Alkyl-quinolines can be found in either enol or keto forms (13). Our structural elucidation indicated that **VI** and MHQ were both in the enol form with the furthest downfield proton corresponding to the hydroxyl proton at 11.6 ppm (Fig. S1-3, Table S3).

To determine if MHQ indeed has dispersal activity, as predicted by the bioactivity-guided fractionation, we tested the activity of the commercial MHQ standard at different concentrations using the DISPEL assay. HPLC-MS indicated that the final active sub-fraction (**VI**) contained roughly 7.3 mM MHQ (Fig. 3A). Purified MHQ was capable of dispersing *P. aeruginosa* at similar levels (Fig. 3B). A dilution series of MHQ revealed that its effective concentration for dispersal activity (EC_50_) is roughly 1 mM (Fig. 3B). Together these data confirmed that MHQ is both made by *P. aeruginosa* and capable of dispersing these bacteria from a surface.

**Figure 3.**
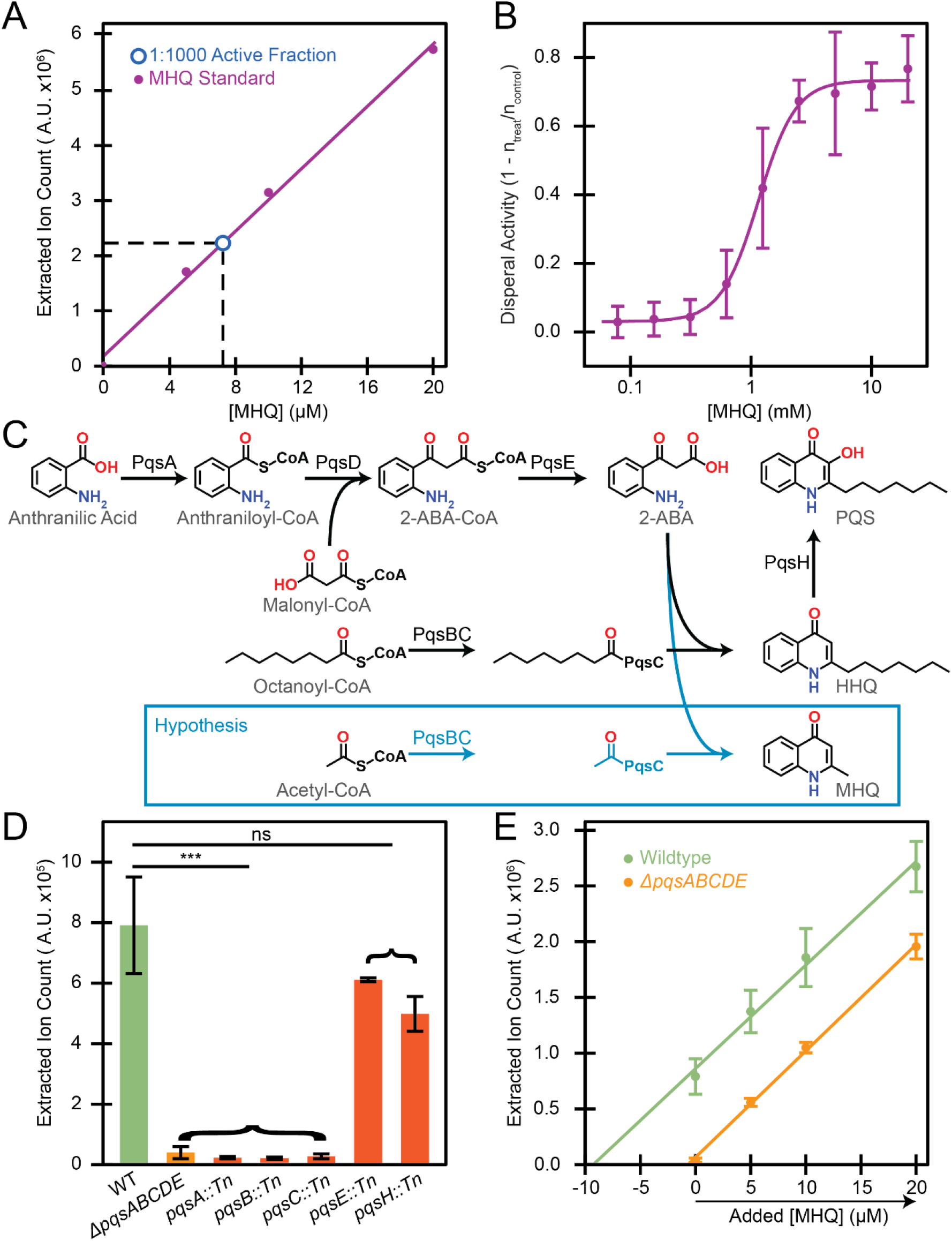
Potency and biosynthesis of MHQ. (A) Quantification of MHQ concentration in **VI** (by HPLC-MS) using commercially available MHQ as a standard. (B) Concentration dependent dispersal activity of MHQ on mid-log (OD_600_ 0.6-0.8) *P. aeruginosa* cells. Mean and standard deviation shown from 4 biological replicates. Fit is based on modified Hill equation (Equation S2). (C) Schematic of the HHQ biosynthetic pathway. Due to the similarity of MHQ to HHQ we hypothesized that it is produced by the same pathway. For consistency with PQS literature MHQ is shown in the keto form. (D) Relative concentration of MHQ in various conditioned media. Mean and Standard deviation shown from 5 biological replicates (WT and *ΔpqsABCDE*) or 2 biological replicates (transposon disruptions). ns p-value > 0.05; *** p-value < 0.001 from Student’s t-test compared to WT conditioned media. (E) An absolute concentration measurement of MHQ in conditioned media (by HPLC-MS) using the standard addition method. Mean and standard deviation shown from 5 biological replicates.

### MHQ is synthesized by the PQS pathway

MHQ is an alkyl quinolone and resembles 2-heptyl-4-hydroxyquinoline (HHQ) with the heptyl tail replaced by a single methyl tail (Fig. 3C). Since HHQ and other alkyl quinolones are synthesized by the enzymes encoded by the *pqsABCDE* operon, we hypothesized that these enzymes may also be responsible for the production of MHQ (Fig. 3C). Specifically, if PqsBC uses acetyl-CoA as a substrate instead of octanoyl-CoA, this would convert the HHQ precursor, 2-ABA, into MHQ (Fig. 3C). To test this hypothesis, we deleted the entire *pqsABCDE* operon and compared the relative amount of MHQ in supernatants from this operon deletion to that of WT (see methods for details). We found that the operon mutant eliminated the HPLC-MS peak associated with MHQ, indicating that these genes are indeed required for producing MHQ (Fig. 3D). To attempt to assess the roles of individual genes within the operon we used mutants containing transposon insertions in each gene (14). We found that interrupting *pqsA, pqsB*, or *pqsC* caused a significant decrease in the amount of MHQ produced, while disruption of *pqsE* or *pqsH* did not have a significant impact on MHQ levels (Fig. 3D). These results are consistent with our hypothesis that MHQ production requires the PqsA, PqsB, and PqsC enzymes. Based on its chemical structure and similarity to HHQ we predicted that PqsE would also be required for MHQ synthesis. However, a recent report suggested that another enzyme (TesB) has redundant activity with PqsE in *P. aeruginosa*, which may explain why MHQ is still produced in this mutant (15).

### Conditioned media contains dilute concentrations of MHQ

Relative concentration measurements were sufficient to establish that the PQS enzymes are required for MHQ synthesis but could not address the absolute levels of MHQ found in conditioned media. Given the complex mixture of molecules in conditioned media that could affect ionization efficiency, we used the standard addition approach to quantify absolute MHQ concentrations. Briefly, we added known concentrations of MHQ to the conditioned media and used HPLC-MS to generate a calibration curve. Extrapolating this calibration curve back to the X-intercept established the absolute concentration of MHQ that would be present with no additional standard. Using this approach, we found that WT conditioned media contained roughly 10 μM MHQ (Fig. 3E). Conditioned media from the *ΔpqsABCDE* mutant contained less than 1 μM MHQ, consistent with our relative concentration measurements (Fig. 3E). Since the EC50 of MHQ for *P. aeruginosa* dispersal is roughly 1 mM (Fig. 3B), the significantly lower concentration of MHQ in the WT conditioned media indicates that either MHQ activity is strongly potentiated in WT conditioned media or there are additional factors in WT conditioned media that lead to its dispersal activity.

To directly determine if MHQ is required for WT dispersal activity, we tested the dispersal activity of conditioned media from the *ΔpqsABCDE* mutant that lacks detectable MHQ. Conditioned media from the *ΔpqsABCDE* mutant retained full activity in the DISPEL assay, supporting the conclusion that the WT dispersal activity does not require MHQ (Fig. S5). In addition to MHQ, the PQS pathway produces a wide range of small molecules with downstream effects on quorum sensing, virulence, and secondary metabolism (16). Thus, the full activity from the conditioned media of the *ΔpqsABCDE* mutant indicates that the MHQ-independent dispersal factors in conditioned media are not the product of the PQS pathway or its downstream signaling.

### MHQ dispersal is not due to antibiotic activity or membrane depolarization

Despite the fact that MHQ is not required for WT conditioned media dispersal activity, MHQ possesses activity on its own and thus could prove to be a useful dispersal agent. Consequently, we sought to determine the mechanism by which MHQ causes dispersal, to both better characterize MHQ and determine how additional dispersal factors might function. MHQ chemically resembles other alkyl quinolones that have been associated with a wide range of biological functions including antibiotic activity and membrane depolarization (16–18). To determine if antibiotic activity could explain dispersal, we used the DISPEL assay to compare the effect of MHQ treatment to treatment with known antibiotics of varying mechanisms of action: novobiocin (replication inhibitor), tetracycline (translation inhibitor), trimethoprim (metabolism disruptor), CCCP (membrane polarity disruptor) and gentamicin (translation inhibitor). None of these treatments resulted in as much dispersal as MHQ treatment, and only CCCP produced any significant dispersal activity (Fig. 4A). Furthermore, we confirmed that most of the antibiotics tested affected cell numbers after a 10-minute treatment comparable to that used in the DISPEL assay, but MHQ had no effect on cell numbers in this time period (Fig. 4B, Fig. S6). These results suggest that antibiotic activity is insufficient to cause the dispersal of *P. aeruginosa*.

**Figure 4.**
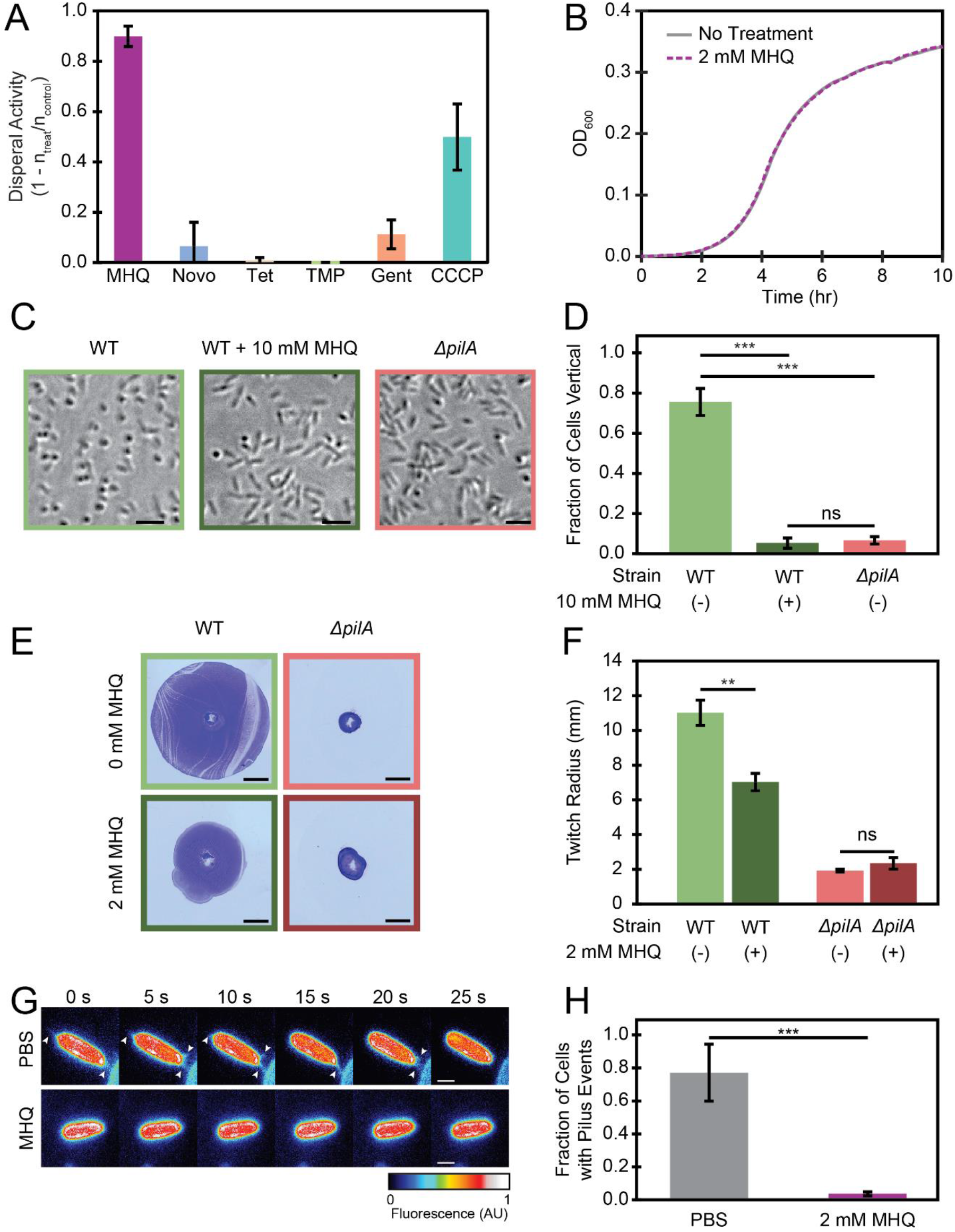
MHQ inhibits Type IV pilus (TFP) activity. (A) Known antibiotics, other than CCCP, do not cause dispersal activity in the DISPEL assay against mid-log (OD_600_ 0.6-0.8) *P. aeruginosa* cells. Mean and standard deviation shown from 3 biological replicates. Novobiocin – 10 mg/ml, Tetracycline – 16 μg/ml, Trimethoprim – 125 μg/ml, Gentamicin – 6 μg/ml, CCCP – 200 μM, MHQ – 2 mM (B) Growth curve of *P. aeruginosa* after 10-minute treatment with MHQ. Mean shown for 3 biological replicates. (C-D) High magnification phase contrast imaging of wildtype *P. aeruginosa* after treatment with PBS (n_total_ = 1347) or MHQ (n_total_ = 1331) for 5 minutes and *ΔpilA* cells treated with PBS (n_total_ = 1995). (C) Scale bar 5 μm. (D) Fraction of cells per image that were aligned vertical to the surface. Mean and Standard deviation shown from 3 biological replicates. ns p-value > 0.05; *** p-value < 0.001 from Student’s t-test. (E-F) TFP dependent twitching assay in the presence or absence of MHQ. (E) Example twitch rings. Scale bar 5 mm. (F) Mean and Standard deviation shown from 3 biological replicates. ns p-value > 0.05; ** p-value < 0.01 from Student’s t-test. (G-H) Microscopic observation of pilus activity treated with PBS (n_total_ = 473) or 2 mM MHQ (n_total_ = 493). (G) Snapshot montage of *P. aeruginosa* cells with labeled pili. Arrowheads indicate extending or retracting pili. Scale bar 1 μm. (H) Fraction of cells per image that had at least one pilus event. Mean and Standard deviation shown from 3 biological replicates. *** p-value < 0.001 from Student’s t-test.

Since CCCP was the only antibiotic that caused even moderate dispersal (though still less than MHQ), and CCCP acts to depolarize bacterial membranes, we sought to determine if MHQ also perturbs membrane integrity. As a quantitative measure of membrane integrity, we used a flow cytometry assay utilizing TO-PRO3, which only stains cells that have been permeabilized, and DiOC_2_(3), which shifts its emission wavelength based on membrane polarization. As positive controls we confirmed that polymyxin-B treatment led to the expected permeabilization of *P. aeruginosa* and CCCP treatment led to the expected depolarization of *E. coli lptD4213* (Fig. S7). We note that we were not able to use DiOC_2_(3) staining in *P. aeruginosa* due to its inability to penetrate the outer membrane, whose integrity is compromised in *E. coli lptD4213*. *P. aeruginosa* cells treated with MHQ did not show an increase in TO-PRO3 staining and *E. coli lptD4213* cells treated with MHQ did not show an increase in DiOC_2_(3) staining. These results indicate that MHQ’s mechanism of action is not mediated by membrane permeabilization or depolarization.

### MHQ inhibits Type IV pilus activity

Since MHQ affects the association of *P. aeruginosa* cells with surfaces, we further investigated its potential mechanism of action by using high-magnification imaging to examine the behavior of individual *P. aeruginosa* cells at the surface. *P. aeruginosa* cells can attach to the surface either by their pole (vertically) or by their side (horizontally) (19–21). Upon treatment with PBS, 76s% (n_total_ = 1347) of the cells attached to the surface vertically. In contrast, upon treatment with 10 mM MHQ, only 5% (n_total_ = 1331) of the cells attached to the surface vertically (Fig. 4C-D).

We noticed a similar behavior between MHQ-treated WT cells and mutants lacking Type IV pilin (TFP) subunits (PilA), as only 7% (n_total_ = 1995) of *ΔpilA* cells were attached vertically even in PBS treatment (Fig. 4C-D). This result suggested that MHQ might disrupt TFP activity. Since TFP are required for twitching motility, we further characterized MHQ’s effect on *P. aeruginosa* in a twitching assay (Fig. 4E). In this assay, cells are placed underneath agar and allowed to spread from their starting spot along the bottom surface of a Petri dish. The extent of this spread is visualized with crystal violet staining and quantified. In the presence of agar made with LB, WT cells traveled 11 +/− 0.7 mm (mean +/− SD) from the starting spot. In contrast, in the presence of agar made with LB and 2 mM MHQ, WT cells traveled significantly less (7 +/− 0.5 mm, Fig. 4F). The effect of MHQ on twitching depended on TFP, as *ΔpilA* cells that lack TFP traveled similar distances away from the starting spot in the absence of MHQ (1.9 +/− 0.1 mm) or in the presence of MHQ (2.3 +/− 0.3 mm) (Fig. 4F).

To directly assay the effect of MHQ on TFP activity we used the recently developed cysteine-labeling approach to fluorescently label TFP and image their dynamics (22–24). This approach uses a cysteine point mutation in an unstructured loop of the PilA pilin subunit to label the pili through maleimide-based click chemistry. When we treated with PBS as a control, we saw TFP extending and retracting from 77% of the cells (n = 473) (Fig. 4G-H and Supplemental Movie S1). In contrast, when cells were treated with 2 mM MHQ for 5 minutes, only 4% (n = 493) of the cells exhibited any TFP extension or retraction events (Fig. 4G-H and Supplemental Movie S2). These results directly demonstrate that MHQ inhibits TFP activity.

## Discussion

Here we sought to disrupt the early stages of surface attachment as a previously unexplored approach to combatting *P. aeruginosa*. We found that *P. aeruginosa* itself produces dispersal agents present in conditioned media. The physiological function for this auto-dispersal activity remains unclear, but there are a variety of high cell density environments where a dispersal agent could be beneficial. For example, in a competition for colonizing a limited surface area, it might be beneficial to prioritize colonization by cells that have managed to grow to higher density by displacing potentially less-fit siblings. Other possibilities include dispersal helping the population respond to conditions of stress by promoting migration away from an undesirable surface or disrupting the surface attachment of competing bacterial species.

Our findings establish MHQ as a small molecule factor that is both produced by *P. aeruginosa* and sufficient to disperse surface-attached *P. aeruginosa.* While MHQ was previously shown to be present in *P. aeruginosa* cells (25), its biosynthesis and functions remained unknown. Here we show that MHQ is synthesized by the *pqsABCDE* operon that is also responsible for the biosynthesis of similar alkyl quinolones like HHQ. Whether MHQ is a byproduct of making HHQ or an intentional product, its characterization presents an opportunity to learn about new ways for dispersing *P. aeruginosa*. Since MHQ was not sufficient to explain the dispersal activity of WT *P. aeruginosa* conditioned media, we are continuing to look for additional factors that either work alone or in conjunction with MHQ to induce dispersal. This work underscores the rich diversity of bioactive secondary metabolites produced by *P. aeruginosa* that have yet to be fully explored and characterized.

Characterizing its mechanism led to the surprising discovery that MHQ inhibits TFP dynamics. TFP inhibition appears to be sufficient to explain MHQ’s activity as MHQ also inhibits TFP-dependent twitching motility and vertical surface attachment. Furthermore, mutants lacking TFP phenocopy MHQ with respect to surface attachment orientation and do not respond to MHQ in a twitching assay. Finally, while MHQ does not affect membrane permeability or polarization, polarization is necessary for TFP activity and CCCP, a known depolarizer, was the only antibiotic capable of producing even mild dispersal. TFP activity is essential for multiple aspects of *P. aeruginosa* colonization and virulence, such that these results suggest that MHQ may be useful for affecting other lifecycle stages beyond early surface attachment. In future studies it will prove interesting to determine the specific mechanism by which MHQ disrupts TFP activity. Others have proposed that disrupting ion channels can inhibit pilus activity (26), but MHQ does not depolarize bacteria. Thus, MHQ may reveal a new way to disrupt TFP and TFP-mediated behaviors.

MHQ has exciting potential applications for both the clinical and lab settings as a small molecule agent that rapidly disperses cells from a surface. MHQ could serve as a therapeutic in the treatment of deadly surface associated pathogens such as *P. aeruginosa.* This could be particularly helpful for combatting the development of biofilms, which are notoriously difficult to treat with conventional antibiotics once formed (27–31). In the future it will also be important to synthesize and characterize MHQ analogs to see if a derivative might increase its potency, as the concentrations currently required for its activity (EC_50_ = 1 mM) may be prohibitively high for some applications. Another potential hurdle is that MHQ’s chemical similarity to other alkyl-quinolones like HHQ could induce crosstalk. We already ruled out some alky-quinolone activities like antibiotic activity and membrane depolarization but establishing the effect of MHQ on other activities like quorum sensing will require further investigation. In any event, our work provides proof-of-principle for future efforts to identify and characterize small molecules that disperse bacteria and disrupt TFP activity.

## Materials and Methods

*P. aeruginosa* PA14 cells were grown 37 °C in liquid LB Miller. The DISPEL Assay was completed by attaching *P. aeruginosa* cells, treating attached cells, washing away dispersed cells, imaging, and using our custom software to determine the number of cells that remained attached. For high-magnification imaging cells were attached to a surface, treated with PBS or MHQ, imaged by both phase contrast and fluorescence. Images were analyzed for their orientation to the surface and extension and retraction events of the TFP. MHQ was purified through an eight-step process involving extraction with resin and ethyl-acetate followed by two open column fractionations and four HPLC fractionations. Additional details on strain growth, strain construction, microscopy, pilus labeling, twitch assays, growth curves, flow cytometry, image analysis, MHQ purification, MHQ structure elucidation, and MHQ quantification by HPLC-MS can be found in the SI Materials and Methods.

## Acknowledgements

NMR was performed in collaboration with Istvan Pelczer (Princeton University NMR Facility). Flow cytometry was performed in collaboration with Christina DeCoste (Princeton University Flow Cytometry Resource Facility [FCRF]). We would like to thank Geoff Vrla for his assistance in strain construction and Jared Balaich for assistance with the HPLC-MS. We appreciate the support and feedback from lab members in the Gitai and Shaevitz labs. Funding was provided in part by NIH (DP1AI124669: Z.G., B.P.B., and R.J.S., T32 GM007388: R.J.S., and, and 1DP2AI124441: M.S.D.). Additional funding provided by the National Science Foundation (NSF PHY-1734030: B.P.B., M.D.K, and C.K.E.), Damon Runyon Cancer Research Foundation (DRG 2385-20: C.K.E.), the German Research Foundation (DFG KO5239/1-1: M.D.K), and for the FCRF by the National Cancer Institute (NCI-CCSG P30CA072720-5921). The opinions, findings, and conclusions or recommendations expressed in this material contents are solely the responsibility of the authors and do not necessarily represent the official views of the NIH or the National Science Foundation.

## Patent Disclosure

A patent describing MHQ as a potential therapeutic in surface dispersal is currently pending.

