## Supplemental Information for "*Pseudomonas aeruginosa* detachment from surfaces via a self-made small molecule"

**This PDF file includes:**

Supplementary Materials and Methods
Figures S1 to S7
Tables S1 to S3
Legends for Movies S1 to S2
SI References

**Other supplementary materials for this manuscript include the following:**

Movies S1 to S2

### Supplemental Materials and Methods

#### *Strains and growth conditions*

The strains can be found in Table S1. We used *Pseudomonas aeruginosa* PA14 for the wildtype strain throughout this study. All strains were grown at 37 °C in liquid LB Miller (Difco).

#### *Strain construction*

The *pilA*-Cys knock-in mutant were generated using two-step allelic exchange (1).Briefly, the cloning vectors were created by digesting the pEXG2 backbone with the HindIII HF restriction enzyme (NEB). Knock-in vectors were created by amplifying the 500 bp flaking regions up- and downstream of the mutation site using primers pilA-T51CC_P1/2/3/4. The overlapping primers pilA-T51C_P2/3 were chosen as reverse complement containing the point mutation. Both flanks were then joined using sewing PCR with the flanking primers. The *pilA*-Cys construct was then ligated into the pEXG2 backbone using T4 DNA ligase (NEB). Next, the cloning vectors were electroporated into *E. Coli* and the correct mutation was confirmed using PCR and sanger sequencing with primers pEXG2_Ver1/2. For mating, 1.5 ml *E. coli* containing the vector were grown to OD 0.5. The *P. aeruginosa* parental strain was grown overnight, and 0.5 ml culture was diluted 1:2 into fresh LB and incubated for 3 hours at 42 °C. Both cultures were concentrated into 100 ml and spotted onto an LB agar plate and incubated overnight at 30 °C. The puddle was scraped off, resuspended into 150 ml PBS, spread onto a VBMM plate containing 30 mg/ml gentamycin and incubated 24 hours at 37 °C. Six single colonies from the VBMM plate were struck onto NSLB and incubated for 24 hours at 30 °C. Several single colonies from the NSLB plate were screened for the correct mutation using PCR amplification with the flaking primers and sanger sequencing.

The *pilA* deletion was constructed by two-step allelic exchange using plasmid pEXG2. Fragments directly upstream and downstream of the *pilA* gene were amplified from gDNA using primer pairs (pilA-1, pilA-2) and (pilA-3, pilA-4), respectively. Upstream and downstream fragments were fused together using overlap-extension PCR with primer pair (pilA-1, pilA-4), and the resulting fragment was cloned into the HindIII site of plasmid pEXG2. The pEXG2 plasmid was integrated into *P. aeruginosa* PA14 through conjugation with the donor strain *E. coli* S17. Exconjugants were selected on 30 µg/mL gentamycin and then the mutants of interest were counter selected on 5% sucrose.

***Microscopy***

*DISPEL assay*

For the DISPEL assay, cells were grown to mid-log ($OD_{600}$ = 0.6-0.8) unless otherwise noted (Fig. 1C) and 50 µl of the culture was added to a well of a 96 well plate (corning black side clear flat bottom). After 10 minutes of attachment cultures were aspirated off slowly. Wells were then incubated for 10 min with 50 µl of treatment. After incubation treatment was aspirated off slowly and the wells were washed gently once with 1x phosphate buffered saline (PBS) and then a final volume of 1xPBS was added for imaging purposes. Wells were imagined on a Nikon TE2000 using a Nikon S Plan Fluor ELWD 20X/0.45 OFN22 PH1, Andor iXon DV-897 and µmanager 2.0 imaging software (2).

*Orientation of individual cells*

For the orientation of individual cells experiments cells were grown to mid-log ($OD_{600}$ = 0.6-0.8) and a 50 µl droplet was placed on a 60 x 22 mm glass cover slip. After 5 minutes of attachment treatment was added to the cells and incubated for 5 minutes. Cells were imaged using a Nikon Ti microscope using a Plan Apo λ 40X PH2, Hamamatsu Orca Flash 4 and Nikon NIS Elements imaging software.

*Pilus labeling*

Pili were labeled as described previously (1, 3, 4) with some modifications. Cells were grown to mid-log growth phase with an $OD_{600}$ of 0.6 and 1 ml of culture was centrifuged at 8000 *x g* for 1 min. The culture supernatant was removed and cell pellets were resuspended in 50 µl of the removed supernatant to concentrate the cells. Concentrated cell suspensions were incubated with 0.5 µl of 0.5 mg/ml stock AF488-malemide (ThermoFisher) for 30 min at room temperature. Cells were centrifuged at 8000 *x g* for 1 min, the supernatant was removed, and 50 µl of the original supernatant was gently added and removed to wash the cells with minimal disturbance to the pellet. The cell pellet was then resuspended in 20-50 µl of the original culture supernatant. 1 µl of labeled cells were added to a 60 x 22 mm glass coverslip and incubated with added PBS as a control or 2 mM MHQ for 5 min. After incubation, a 1% agarose (Invitrogen) PBS pad was used to sandwich cells between the coverslip and pad. 1% agarose PBS pads were made with 2 mM MHQ for imaging MHQ-treated cells. Cells were imaged using a Nikon Ti microscope using a Plan Apo λ 100X Oil PH3, GFPHQ

Filter Cube (Ex:455-485, Em:500-545), Hamamatsu Orca Flash 4 and Nikon NIS Elements imaging software.

***Twitching assay***

Colonies of cells were picked and stabbed through the agar of a plate and placed at 30 °C for 4 days. After 4 days the agar was removed from the dish and 0.5% crystal violet was added to the plate. After 5 minutes of staining the crystal violet was removed and the plate was washed 3 times with water. The stained rings were imaged using a Canon EOS Rebel T1i (Lake Success, NY) and measured in FIJI (5–7).

***Flow cytometry***

Flow Cytometry protocol was adapted from (8). In brief, overnight *E. coli lptD4213* and *P. aeruginosa* PA14 cultures were grown to early-mid exponential phase (OD600 = 0.4-0.6) at 37°C. Each culture was then diluted 1:10 into 1x PBS and treated with either 2 mM MHQ, 5 uM CCCP, 4 µg/mL polymyxin for 10 minutes. Cells were stained with the BacLight Bacterial Membrane Potential kit (ThermoFisher B34950). This kit uses DiOC2(3) to measure a cell's membrane potential as a ratio of green (488 nm ex, 525/50 nm em) to red (488 nm ex, 610/20 nm em) (9). Membrane integrity was measured by staining cells with TO-PRO-3, a dye that is excluded from cells with an intact membrane (640 nm ex, 670/30 nm em). The LSRII flow cytometer (BD Biosciences) at the Flow Cytometry Resource Facility, Princeton University, was used to measure the fluorescent intensities of both dyes in response to antibiotic or MHQ treatment.

100,000 events were recorded for each data file. Data was analyzed using FlowJo v10 software (FlowJo LLC, Ashland, OR).

***Growth curves***

Mid-log ($OD_{600}$ = 0.6-0.8) *P. aeruginosa* cells were treated with antibiotics and MHQ (Novobiocin – 10 mg/ml, Tetracycline – 16 µg/ml, Trimethoprim – 125 µg/ml, Gentamicin – 6 µg/ml, CCCP – 200 µM, MHQ – 2 mM) for 10 minutes. Cells were spun and resuspend in equal volumes of drug free LB twice. Cells were diluted 1:100 and grown for 10 hrs in a microplate reader (Tecan) at 37 °C with shaking.

***Image analysis***

*DISPEL assay*

Images were analyzed using custom Matlab (Mathworks) scripts to count the number of cells in the image. The data was then normalized using the control images of the PBS treated or LB treated cells depending on the similarity to the treatment. Activity was defined as

$$Activity = 1 - \frac{Average\ \#\ of\ cells\ in\ treatment\ well}{Average\ \#\ of\ cells\ in\ control\ well} \qquad \text{(S1)}$$

Data was fit based on using a modified Hill equation

$$y = a * \frac{x^n}{{EC_{50}}^n + x^n} + b \qquad \text{(S2)}$$

with *y* being the activity and *x* being the condition varied in the experiment. *a* and *b* sum to 1 and are the relative magnitude of the experimental variant and the no treatment offset, respectively. $EC_{50}$ is the effective concentration at which the activity is 50% of the total effect. The cooperativity coefficient, *n*, refers the steepness of the transition between effect and no effect. For our data *n* was around 15.

*Orientation of individual cells*

Individual cells were hand-scored for their orientation to the surface as either vertical or horizontal. For each of 3 biological replicates an image containing 250-1000 cells was scored. The fraction of cells vertical was calculated by dividing the number of cells vertical by the total number of cells scored. The average and standard deviation across biological replicates is shown (Fig. 4B).

*Pilus activity*

Individual cells were hand-scored for whether there was a pilus event within a 5-minute movie. A pilus event was generously defined as any extension or retraction event regardless if the cycle was fully completed during the movie. For each of 3 biological replicates an image containing 50-250 cells was scored. The fraction of cells with pilus activity was calculated by dividing the number of cells with a pilus event by the total number of cells scored. The average and standard deviation across biological replicates is shown (Fig. 4F).

***Large scale culture and fractionation***

For the large-scale culturing and purification of MHQ, 1 ml overnight culture was used as an inoculum for each 500 ml LB media batch and cultivated overnight, at 37 °C shaken at 200 rpm. A total of 50 L total was generated. 2 batches at a time were combined and centrifuged at 15,000 xg for 15 min (JLA 9.1000, Avanti JXN-30, 25°C) to separate cells from the conditioned media. The conditioned media was shaken with Diaion HP-20 (Sigma-Aldrich), 50 ml for 1L conditioned media, for 1h and resins were eluted with MeOH. Subsequently, the extracts from all batches were combined and dried in vacuo. The total extract was re-suspended with 500 ml $H_2O$, then partitioned with 500 ml of ethyl acetate 15 times. The organic layers from each partition were combined and dried in vacuo

The dried crude extract was re-suspended in 5 ml MeOH and subjected to open column chromatography by using Mega Bond Elut-C18, 70 g, 150 ml (Agilent Technologies, USA) with stepwise elution with MeCN : $H_2O$ (10:90, 20:80, 20:80, 30:70, 30:70, 50:50) and a final wash 100% MeOH. At this point and each subsequent step of the purification process a portion of each fraction was resuspended in 1x PBS and used in the DISPEL assay to determine activity (Fig. 2A-B and Methods: DISPEL assay).

Active fraction was further purified on another Mega Bond Elut-C18, 10 ml (Agilent Technologies, USA) with stepwise elution with MeCN: $H_2O$ (10:90**,** 50:50**,** 100:0) and a final wash 100% MeOH.

Active fraction was purified by HPLC on a C18 reverse phase column (Poroshell 120 EC-C18, 9.4 x 250 mm, 4 µm) with gradient 0-15 min: 30- 100 % A, 15- 21 min: 100 % A, 21- 21.5 min: 100- 30 % A, at a flow rate of 1.25 ml/min. (Solvent A: MeCN with 0.01% TFA; B: $H_2O$ with 0.01% TFA).

The active fraction was further purified by HPLC on a C18 reverse phase column (Poroshell 120 EC-C18, 9.4 x 250 mm, 4 µm) with gradient 0-12 min: 20-60 % A, 12- 12.1 min: 100 % A, 12.1- 16.8 min: 100 % A, 16.8- 17 min: 100- 20

% A, at a flow rate of 1.25 ml/min. (Solvent A: MeCN with 0.01% TFA; B: $H_2O$ with 0.01% TFA).

The active fraction was further purified by HPLC on a C18 reverse phase column (Poroshell 120 EC-C18, 9.4 x 250 mm, 4 µm) with isocratic elution 0-23 min: 15 % A, at a flow rate of 1.3 ml/min. (Solvent A: MeCN with 0.01% TFA; B: $H_2O$ with 0.01% TFA).

The active fraction was further purified by HPLC on a C18 reverse phase column (Poroshell 120 EC-C18, 4.6 x 250 mm, 4 µm) with isocratic elution 0- 17 min: 17 % A, at a flow rate of 0.6 ml/min. (Solvent A: MeCN with 0.01% TFA; B: $H_2O$ with 0.01% TFA). Finally, 1.3 mg (0.09 mg/ L) of final fraction was obtained (**VI,** Fig. 2B).

***Structural elucidation of MHQ***

**VI** was isolated as a dark orange solid. HPLC-HRMS (using a Shimadzu HPLC and Thermo LTQ Orbitrap XL MS) established that the majority of **VI** corresponded to the *m/z* = 160.07559 $[M+H]^+$, with a predicted molecular formula $C_{10}H_9NO$ (calculated m/z = 160.0762). $^1$H-NMR spectrum of **VI** indicated one hydroxyl, four aromatic, one methine, three methyl protons (Table S3). In the $^{13}$C-NMR spectrum, ten unique signals were observed, including nine aromatic carbons, one methyl (Table S3). The bicyclic structure of 2-methyl-4-hydroxyquinoline (MHQ) seen in Fig. 2E was elucidated by COSY and HMBC experiments (Fig. S1-3). The structure of MHQ was further confirmed by comparison with MS, $^1$H-NMR and $^{13}$C-NMR spectra for a commercially available authentic standard of the same molecule (Sigma) as well as $^1$H-NMR data for

MHQ from a previous reference reporting the same molecule (Fig. S3 and Table S4) (10).

***HPLC-MS curve information***

For both relative and absolute quantification of MHQ in conditioned media and **VI** we used HPLC-MS (Agilent Single Quad) using a C18 reverse phase column (Poroshell 120 EC-C18, 4.6 x 100 mm, 2.7 µm). For standard addition, the desired concentration of MHQ was added to the sample and then run on the HPLC-MS. MHQ ion count was determined by integrating the peak of the extracted ion chromatogram for 160.1. The absolute concentration of MHQ was calculated for the conditioned media by taking the intercept of the linear fit for the standard curve in conditioned media.

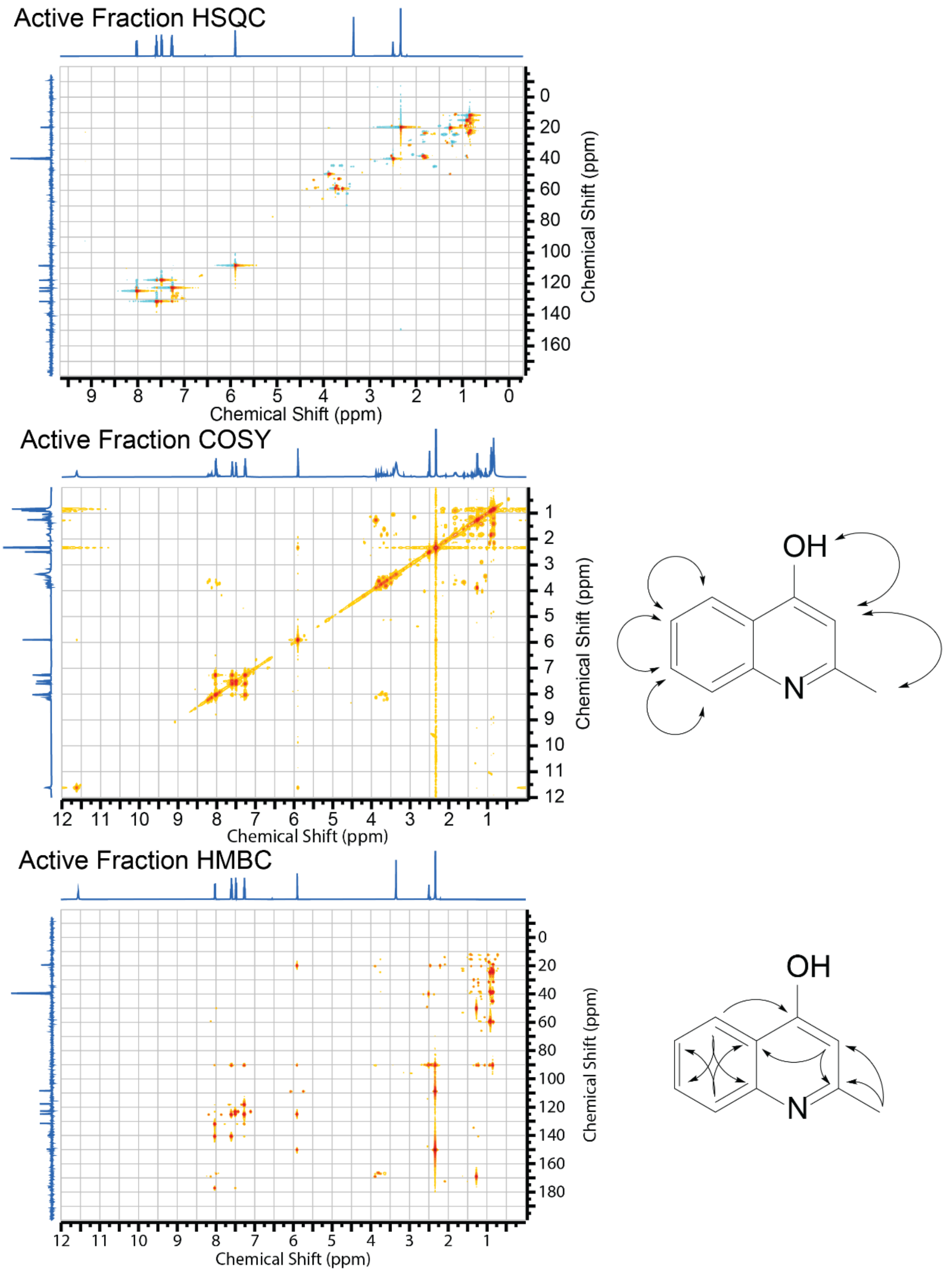


**Fig. S1.** 2D NMR spectra (HSQC, COSY, HMBC) of **VI** used to elucidate the structure of MHQ. All spectra were measured in DMSO-d6 at 295 K. The structure of MHQ is shown with selected correlations full arrows (COSY) and half arrows (HBMC).

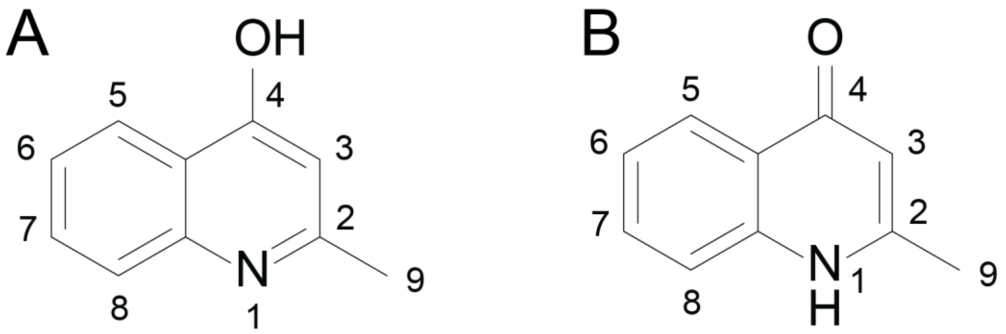


**Fig. S2.** Tautomers of MHQ shown in enol (A) and keto (B) forms. Nuclear numbering for peak assignment in Table S3.

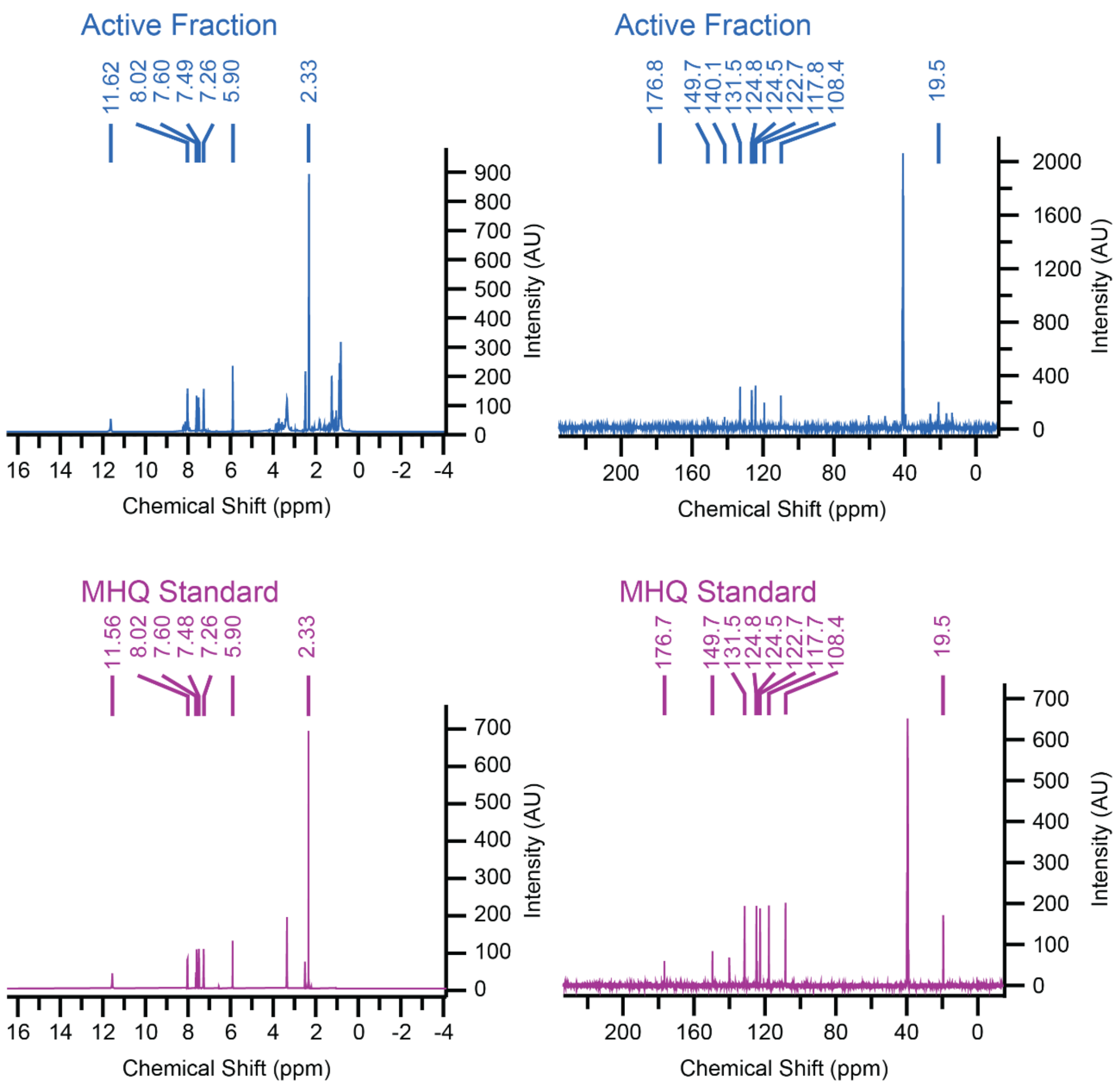


**Fig. S3.** 1D NMR spectra ($^{1}$H-NMR and $^{13}$C-NMR) of **VI** and commercially available MHQ. All spectra were measured in DMSO-d6 at 295 K.

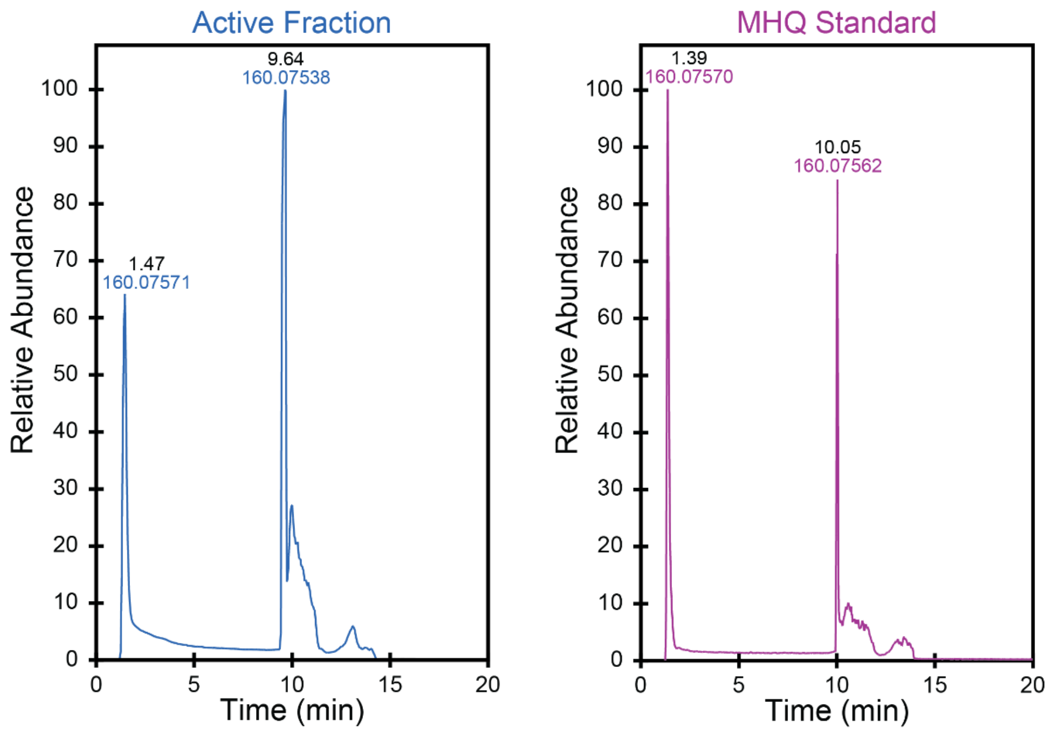


**Fig. S4.** HPLC-HRMS retention time traces of 160.07559 m/z in **VI** and commercially available MHQ.

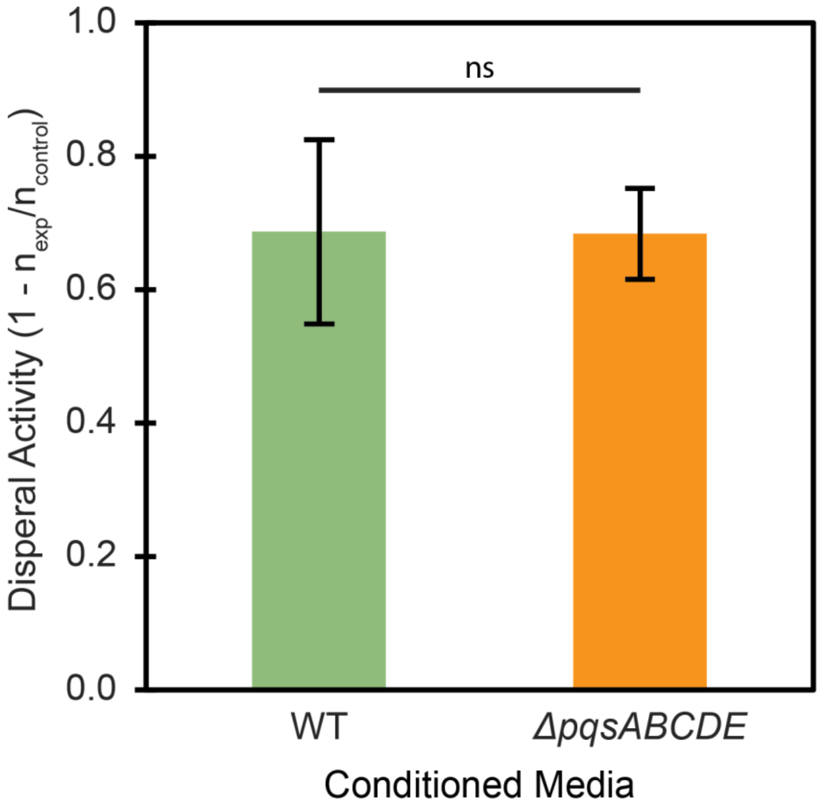


**Fig. S5.** Conditioned media from the *ΔpqsABCDE* has full dispersal activity. Mean and standard deviation shown from 5 biological replicates. ns p-value > 0.05 from Student's t-test**.**

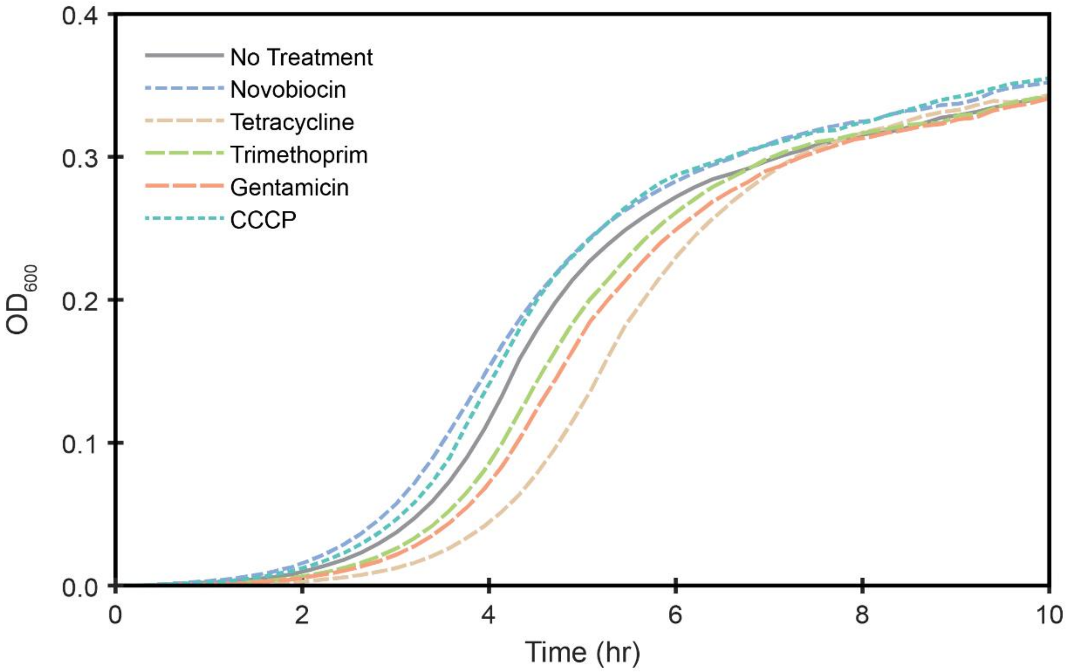


**Fig. S6.** Growth curves of *P. aeruginosa* after treatment with known antibiotics for 10 minutes. Mean shown for 3 biological replicates.

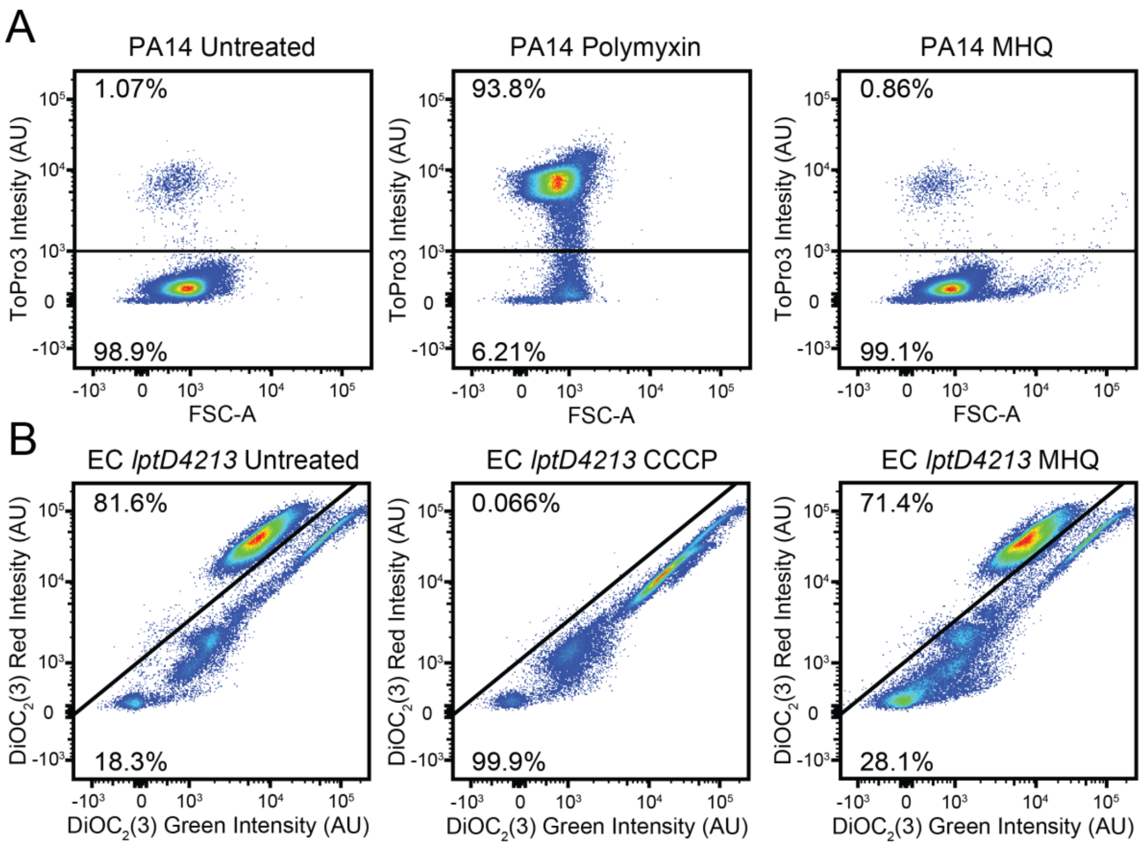


**Fig. S7.** Flow Cytometry analysis of membrane integrity and depolarization following MHQ treatment. (A) *P. aeruginosa* cells were stained with ToPro3 following either no treatment, polymyxin treatment, or MHQ treatment. Increased ToPro3 staining is indicative of cell permeabilization. (B) *E. coli lptD4213* cells were stained with DiOC2(3) following either no treatment, CCCP treatment, or MHQ treatment. Fluorescence shift towards green indicate membrane depolarization.

**Table S1.** Bacterial strains

| **Strain description** | **Unique identifier** | **Reference** |
|---|---|---|
| *P. aeruginosa strain UCBPP PA14* | ZG 38 | Lab stock |
| PA14 *ΔpilA* | ZG 1713 | This study |
| PA14 *ΔpqsABCDE* | ZG 1714 | (11) |
| PA14 *pqsA::MAR2xT7* | 23621 | (12) |
| PA14 *pqsB::MAR2xT7* | 42596 | (12) |
| PA14 *pqsC::MAR2xT7* | 32423 | (12) |
| PA14 *pqsE::MAR2xT7* | 45262 | (12) |
| PA14 *pqsH::MAR2xT7* | 47950 | (12) |
| PA14 *pilA-T51C* | ZG 1715 | This study |
| *E. coli lptD 4213* | ZG 1598 | Lab stock |

**Table S2.** Primers

| Primer name | Primer sequence | Usage |
|---|---|---|
| pilA-1 | GAT ACA AAG CTT CTT GTT GCG CTG GGC CTG | *pilA* deletion |
| pilA-2 | GGT ACC TGC AGT CAG GGC CGC AAC CAC GAT CAT CAG TTC | *pilA* deletion |
| pilA-3 | GAA CTG ATG ATC GTG GTT GCG GCC CTG ACT GCA GGT ACC | *pilA* deletion |
| pilA-4 | GAT ACA AAG CTT CAT GAA CAA GAG CAA GCG GC | *pilA* deletion |
| pilA-T51CC_P1 | GATACAAAGCTTCCGCTGAGTTGAATTGTGTCG | *pilA* Cysteine Replacement |
| pilA-T51CC_P2 | CTGGCTGCCAGCGCCAAGTGTCTTATTGGCGATAGCTCTGCC | *pilA* Cysteine Replacement |
| pilA-T51CC_P3 | GGCAGAGCTATCGCCAATAAGACACTTGGCGCTGGCAGCCAG | *pilA* Cysteine Replacement |
| pilA-T51CC_P4 | GATACAAAGCTTCCACCACAAACAGATGATTGCC | *pilA* Cysteine Replacement |
| pEXG2_Ver1 | GTTGCATGGGCATAAAGTTGCC | Confirming inserts |
| pEXG2_Ver2 | CGGGTCCTCAACGACAGG | Confirming inserts |
| PA14pilA_Seq1 | GGCTGTTCAGGTCGCAGTAGG | Confirming Exconjugants |

**Table S3.** NMR peak assignments

| | Final Active Fraction (VI) | | MHQ Standard | | Pseudan I* (10) |
|---|---|---|---|---|---|
| No. | $^{1}H$ δ (*J*, Hz) (500.40 MHz) | $^{13}C$ δ (125.84 MHz) | $^{1}H$ δ (*J*, Hz) (500.40 MHz) | $^{13}C$ δ (125.84 MHz) | $^{1}H$ δ (*J*, Hz) |
| 2 | | 149.7 | | 149.6 | |
| 3 | 5.90 (s) | 108.4 | 5.90 (s) | 108.4 | 6.01 (s) |
| 4 | | 176.8 | | 176.7 | |
| | 11.62 (s, OH) | | 11.56 (s, OH) | | 11.35 (br s, OH) |
| 4a | | 124.5 | | 124.5 | |
| 5 | 8.02 (d, 9.5) | 124.8 | 8.02 (d, 9.6) | 124.8 | 8.16 (d) |
| 6 | 7.26 (t, 7.5) | 122.7 | 7.26 (t, 8.1) | 122.7 | 7.21 (t) |
| 7 | 7.60 (t, 8.4) | 131.5 | 7.60 (t, 8.4) | 131.5 | 7.53 (m) |
| 8 | 7.49 (d, 8.3) | 117.8 | 7.48 (d, 8.2) | 117.7 | 7.48 (m) |
| 8a | | 140.1 | | 140.1 | |
| 9 | 2.33 (s) | 19.5 | 2.33 | 19.5 | 2.34 (s) |

*Pseudan I was the name given to MHQ in (10)

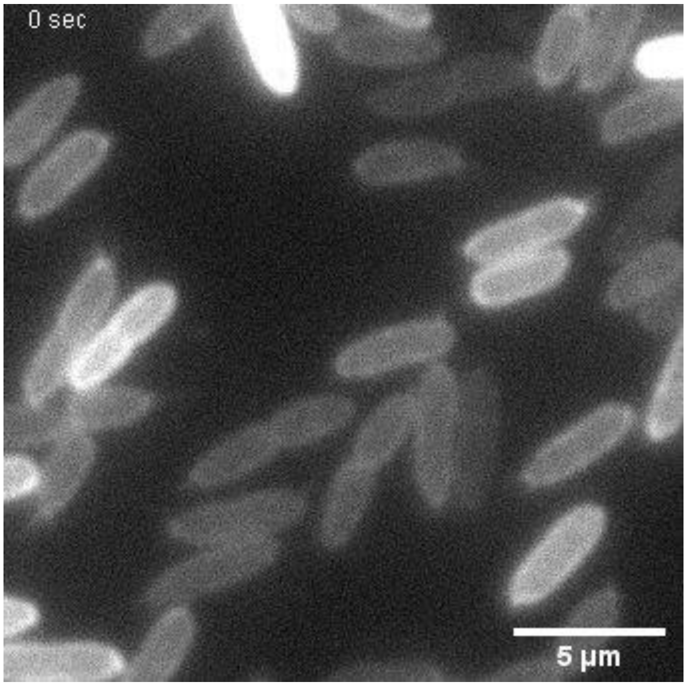


**Movie S1 (separate file).** Pilus activity of PBS Treated *P. aeruginosa* cells. Pili are fluorescently labeled. To account for photobleaching images were normalized to saturate 0.3% of pixels in each frame.

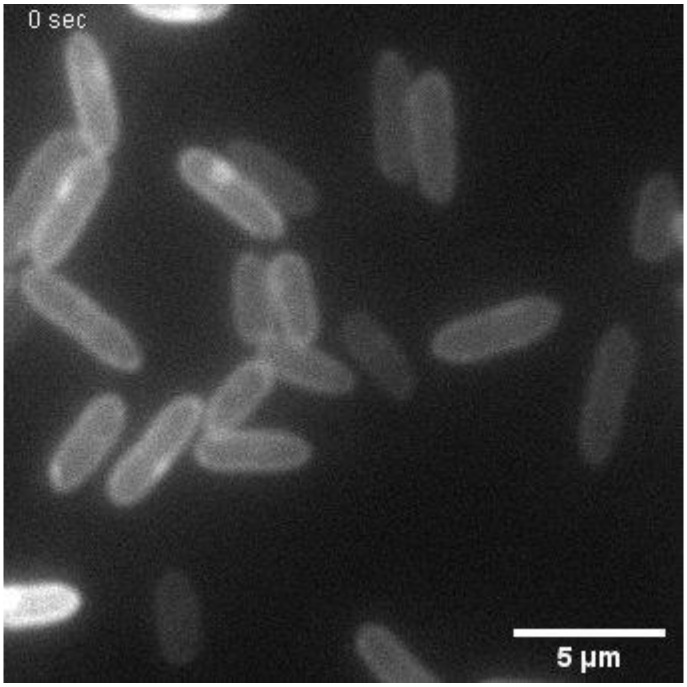


**Movie S2 (separate file).** Pilus activity of MHQ Treated *P. aeruginosa* cells. Pili are fluorescently labeled. To account for photobleaching images were normalized to saturate 0.3% of pixels in each frame.
